# Knockout of the gene encoding the extracellular matrix protein SNED1 results in early neonatal lethality and craniofacial malformations

**DOI:** 10.1101/440081

**Authors:** Anna Barqué, Kyleen Jan, Emanuel De La Fuente, Christina L. Nicholas, Richard O. Hynes, Alexandra Naba

## Abstract

**Background:** The extracellular matrix (ECM) is a fundamental component of multicellular organisms that orchestrates developmental processes and controls cell and tissue organization. We previously identified the novel ECM protein SNED1 as a promoter of breast cancer metastasis and showed that its level of expression negatively correlated with survival of breast cancer patients. Here we sought to identify the roles of SNED1 during murine development and in physiology.

**Results:** We generated two novel *Sned1* knockout mouse strains and showed that *Sned1* is essential since homozygous ablation of the gene led to ~67% early neonatal lethality. Phenotypic analysis of the surviving knockout mice obtained revealed a role for SNED1 in the development of craniofacial and skeletal structures since *Sned1* knockout resulted in growth defects, nasal cavity occlusion, and craniofacial malformations. *Sned1* is widely expressed in embryos, notably in neural-crest derivatives. We further show here that mice with a neural-crest-cell-specific deletion of *Sned1* survive, but display facial anomalies partly phenocopying the global knockout mice.

**Conclusions:** Our results demonstrate requisite roles for SNED1 during development and neonatal survival. Importantly, the deletion of 2q37.3 in humans, a region that includes the *SNED1* locus, has been associated with facial dysmorphism and short stature.

## Introduction

The extracellular matrix (ECM) is a fundamental component of multicellular organisms. It is a complex and dynamic assembly of hundreds of proteins that provides support to cells and instructs their behaviors.^1–3^ Mutations in ECM genes or changes in ECM composition, architecture or abundance have been shown to cause or accompany a plethora of diseases including fibrosis, renal, skeletal and vascular diseases, and cancers.^3–6^ Knockout mouse models have allowed the identification of the instrumental roles of ECM proteins and ECM-protein receptors at multiple steps of embryonic development^7–15^ Knockout of several sub-units of the ECM receptors of the integrin family causes early embryonic lethality.^16,17^ More specifically, and of relevance to the present study, ECM proteins including collagen, fibronectin, tenascin-C, versican, and aggrecan have been shown to regulate important features such as the migration of neural crest cells (NCCs).^18–21^

The murine gene encoding the ECM protein SNED1 was cloned by Leimeister and collaborators in 2004 from renal stromal cells.^22^ In this original publication, the authors used *in-situ* hybridization and showed that *Sned1* is broadly expressed throughout development, mainly in the ectoderm, neural-crest, and mesoderm derivatives including kidneys, adrenal glands, lungs, skeleton, limbs, and the head region. Yet, the functions of SNED1 are, until now, unknown. In a proteomic screen comparing the ECM composition of poorly and highly metastatic human mammary tumors, we identified SNED1 as being more abundant in more aggressive mammary tumors.^5^ We further showed that SNED1 functioned as a metastatic promoter, since knocking down *SNED1’s* expression in tumor cells prevented metastasis.^5^ Importantly, we also demonstrated that *SNED1* expression level was predictive of survival of hormone-receptor negative breast cancer patients.^5^ Additionally, *Sned1* was found to be up-regulated in murine pancreatic ductal adenocarcinoma (PDAC) cells cultured in a 3-dimensional ECM and which are more resistant to chemotherapeutic drugs, as compared to cells grown on a 2dimensional substrate.^23^ *Sned1* knockdown was also shown to inhibit mutant-p53-driven murine PDAC cell invasion *in vitro*^24^ While SNED1 has been shown to regulate cancer cell phenotypes, the roles of this protein in embryonic development and physiology remain to be identified.

To address this question, we generated knockouts of *Sned1* in mice. Using these novel mouse models, we demonstrate here that *Sned1* is an essential gene, since *Sned1* knockout causes ~67% early neonatal lethality within p0 and p2. Over the course of our study, some homozygous knockout mice survived. Interestingly, they were easily distinguishable from wild-type and heterozygous littermates because of their smaller size and craniofacial malformations. Considering the craniofacial phenotype observed in knockout mice and the pattern of expression of *Sned1*, we further sought to examine the potential role of SNED1 on the NCC population, which contributes to form most of the bones and cartilage of the head.^2,5–28^ We thus genetically ablated *Sned1* in the NCCs and observed that while *Sned1^NCC-/NCC-^* mice survived, they presented growth defects and facial anomalies resembling malformations observed in the global knockout animals. Altogether, our results point to a role for SNED1 in the proper formation of craniofacial structures, and that altered craniofacial structures resulting from *Sned1* knockout may be an explanation for the early neonatal lethality of *Sned1* knockout mice.

## Results

### *Sned1* is conserved among all vertebrates and presents key features of ECM proteins

The *SNED1* (<u>S</u>ushi, <u>N</u>idogen and <u>E</u>GF-like <u>D</u>omains <u>1</u>) gene, initially named *SNEP* (<u>S</u>tromal <u>N</u>idogen <u>E</u>xtracellular matrix <u>P</u>rotein), encodes a secreted protein composed of characteristic domains commonly found in ECM proteins^29,30^ including an amino-terminal NIDO domain, 15 EGF-like and calcium-binding EGF-like domains, one or two Follistatin-N-terminal-like (FOLN) domains (depending on the prediction algorithms^31,32^), a Complement Control Protein (CCP, also known as Sushi domain), and three Type III fibronectin (FN3) domains in the carboxy-terminal region of the protein (Figure 1). Interrogation of gene and protein databases revealed that orthologues of *SNED1* are found in all sequenced vertebrates, including fish, reptiles, amphibians, birds, and mammals (Figure 1), but not in lower organisms. Of note, the evolution of vertebrates and mammals has been accompanied by the expansion of families of ECM proteins existing in lower organisms (such as the collagens) and by the appearance of novel ECM proteins,^33–35^ such as SNED1. This increasing complexity of the ECM has been concomitant with the appearance of novel structures including the neural crest and endothelium-lined vasculature in vertebrates or of mammary glands in mammals.^36,37^

**Figure 1.**
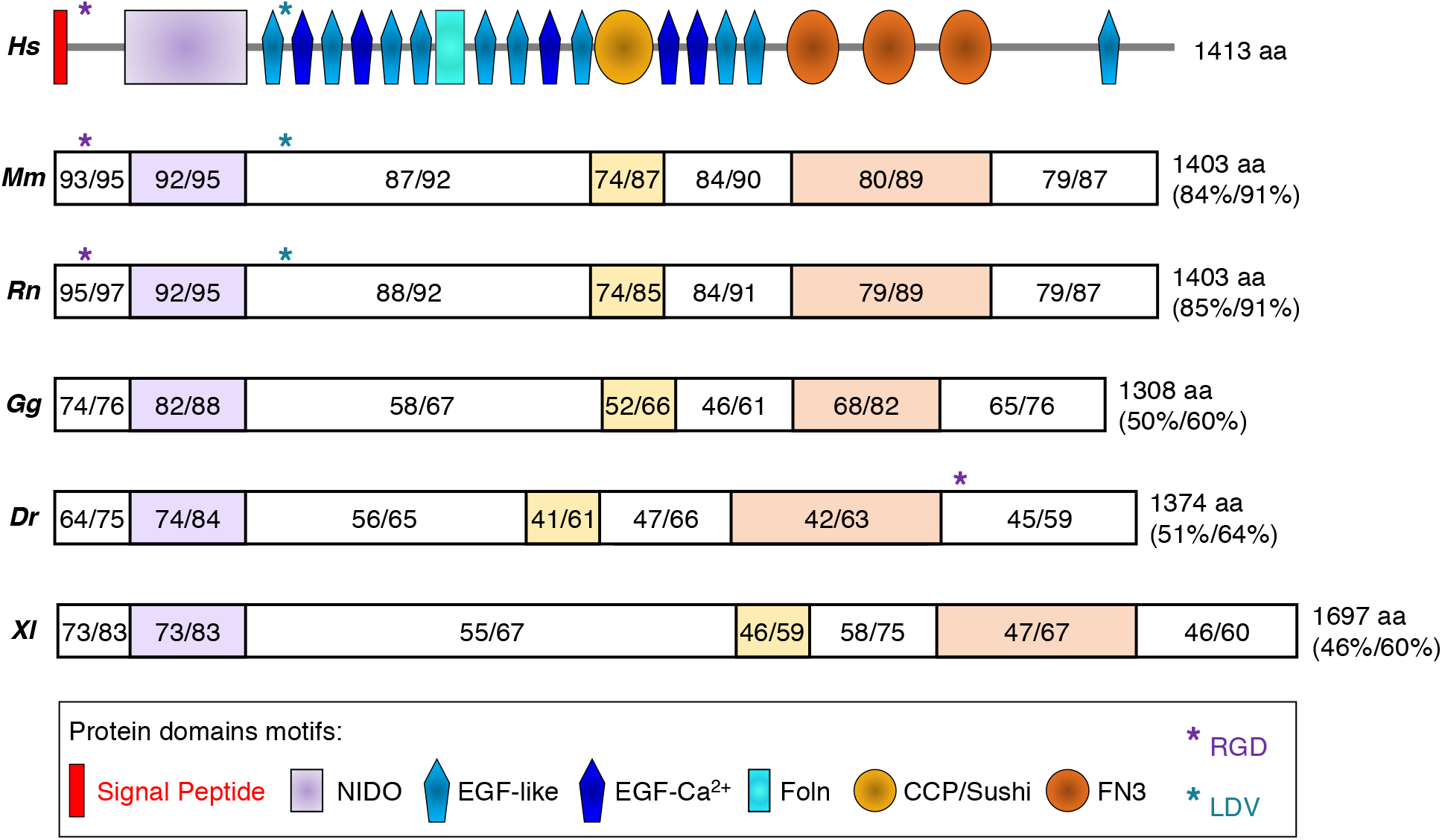
SNED1 phylogeny. Schematic representation of the protein domains of SNED1 and alignment between the human (*Hs*) sequence of SNED1 (UniProt accession Q8TER0) and orthologs found in mouse (*M. musculus, Mm*; UniProt accession Q70E20); rat (*R. Norvegicus, Rn*; UniProt accession Q5ZQU0); chicken (*G. gallus, Gg*; UniProt accession A0A1D5P671); zebrafish (*D. rerio, Dr*; UniProt accession E7F2S5); and frog (*X. laevis, Xl*; UniProt accession A0A1L8GAZ2). Alignment was performed using Protein BLAST (https://blast.ncbi.nlm.nih.gov/Blast.cgi?PAGE=Proteins). Percent identity and percent homology are indicated for each domain or fragment of SNED1 and for the full-length proteins. Protein length in amino acid (aa) is also indicated.

In addition, sequence analysis has revealed the presence of an RGD motif in the carboxy-terminal region of the zebrafish sequence and two consensus potential binding sites, RGD and LDV, for integrins,^38,39^ which are known ECM protein receptors, in the amino-terminal region of mammalian sequences of SNED1 (Figure 1), suggesting that the protein has further evolved, likely to support the development of mammalian-specific structures, as has occurred for other ECM proteins.

### *Sned1* knockout results in early neonatal lethality

*Sned1* was targeted using the “knockout-first allele” construct designed through an effort supported by the Wellcome Trust Sanger Institute and the Knockout Mouse Project^40^. In a first approach, *Sned1* knockout was achieved using an IRES:LacZ-trapping cassette^40^ inserted in the second intron of *Sned1* (Figure 2A). Decreased *Sned1* mRNA levels were confirmed using RT-qPCR on RNA extracted from mouse embryonic fibroblasts generated from *Sned1^WT/WT^, Sned1^WT/LacZ-Neo^, Sned1^LacZ-Neo/LacZ-Neo^* mice (Figure 2C).

**Figure 2.**
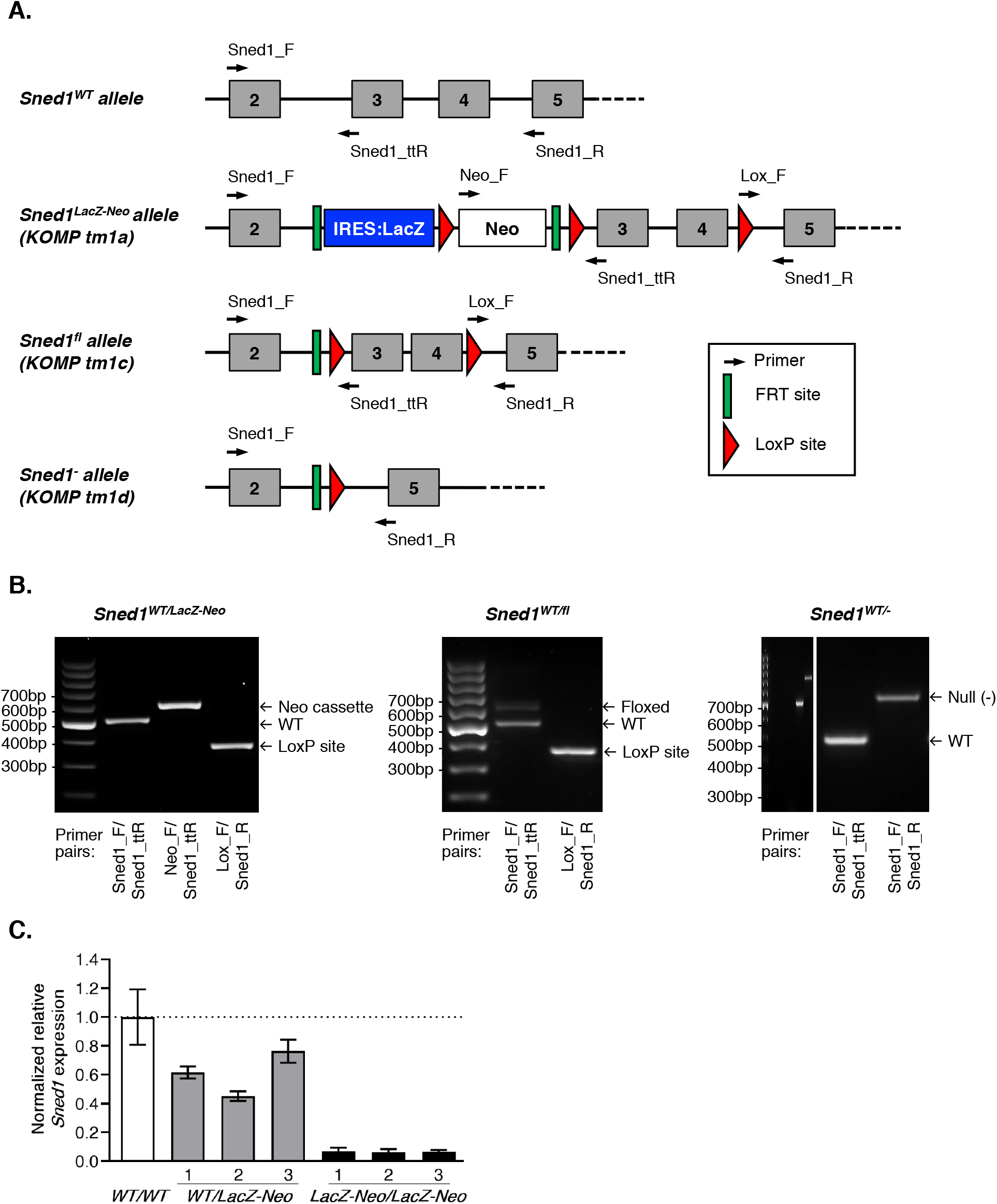
*Sned1* knockout strategies and genotyping. **A.** Schematics of the different alleles *Sned1^WT^, Sned1^LacZ-Neo^* (*KOMP tm1a*), *Sned1^fl^* (*KOMP tm1c*), and *Sned1^-^* (*KOMP tm1d*). Arrows indicate primers used for genotyping. **B.** PCR genotyping of genomic DNA isolated from tail samples of *Sned1^WT/LacZ-Neo^* (left panel), *Sned1^WT/fl^* (middle panel), and *Sned1^WT/-^* (right panel) mice. **C.** RT-qPCR was used to monitor the level of expression of *Sned1* in mouse embryonic fibroblasts isolated from *Sned1^WT/WT^* (n=1), *Sned1^WT/LacZ-Neo^* (n=3), or *Sned1^LacZ-Neo/LacZ-Neo^* (n=3) embryos. Bar charts represent normalized expression of *Sned1* relative to wildtype cells. Actin expression level was used to normalize RT-qPCR data. Data are presented as means ± S.E.M.

We intercrossed heterozygous (*Sned1^WT/LacZ-Nea^*) mice and examined 55 litters born from 16 different breeding pairs, representing a total of 334 pups, of which 202 were still alive at weaning. We observed that a significant number of pups died within 48 hours after birth, and, when carcasses could be retrieved for genotyping, we found that the majority of dead neonates were knockout (*Sned1^LacZ-Neo/LacZ-Neo^*) animals, although wild-type and heterozygous pups were also found dead and contribute to explain the overall 60% viability observed. We further observed that at weaning (p21 +/- 1 day), knockout mice were present in sub-Mendelian ratio (8.9% instead of the expected 25%; χ^2^=20.303; two-tailed p<0.0001) indicating a 33% viability of the knockout pups (Table 1). However, Mendelian ratios were observed for E18.5 embryos obtained by C-section (6 litters analyzed, χ^2^=1; two-tailed p=0.607), suggesting that lethality of knockout pups occurred either around or shortly after birth (Table 1).

**Table 1.** Numbers of expected and observed E18.5 embryos and 21-day-old pups of each genotype obtained by crossing heterozygous *Sned1^WT/LacZ-Neo^* mice.

| *Sned1*<sup>WT/LacZ-Neo</sup> x<br>*Sned1*<sup>WT/LacZ-Neo</sup> | *Sned1*<sup>WT/WT</sup> | *Sned1*<sup>WT/LacZ-Neo</sup> | *Sned1*<sup>LacZ-Neo/LacZ-Neo</sup> | $\chi^2$<br>p-value |
| --- | --- | --- | --- | --- |
| **Expected** | 25% | 50% | 25% |  |
| **E18.5** | 14 (27.5%) | 22 (43.1%) | 15 (29.4%) | 0.607 |
| **Alive at weaning (p21)** | 54 (26.7%) | 130 (64.4%) | **18 (8.9%)** | **<0.0001** |

We then generated a conditional knockout mouse strain by crossing heterozygous *Sned1^WT/LacZ-Neo^* mice with mice expressing the Flp recombinase under the control of the actin promoter. This resulted in the excision of the IRES:LacZ and Neo cassettes, and the generation of a floxed allele (*Sned1^fl^*, Figures 2A and 2B). *Sned1^fl/fl^* mice were indistinguishable from wild-type mice indicating that the excision of the gene-trap cassette reverted the phenotype. This strain will be useful to determine the function of *Sned1* in a cell- or time-specific manner (*see below*). We further crossed conditional knockout mice with mice expressing the Cre recombinase under the control of a CMV promoter, this introduces a frameshift mutation that triggers nonsense-mediated decay of the truncated transcripts^40^ and ultimately generates a null allele (*Sned1*^-^, Figures 2A and 2B). Heterozygous mice (*Sned1^WT/-^*) were further intercrossed to generate homozygous null mice (*Sned1*^-/-^). We evaluated 13 litters born from 7 different breeding pairs of heterozygous mice (representing a total of 88 pups alive upon inspection of the cages on the expected date of birth, of which 63 were still alive at weaning). Similarly to what we observed with the gene-trap allele, a significant number of pups died within 48 hours after birth. When carcasses were retrieved for genotyping, we found that the majority of dead neonates were *Sned1*^-/-^ mice. We further observed that at weaning (p21 +/- 1 day), null mice (*Sned1*^-/-^) were present in sub-Mendelian ratio (7.9%, instead of the expected 25%; χ^2^=12.492; two-tailed p=0.0019), indicating that null pups had died before that (Table 2).

**Table 2.** Numbers of expected and observed E18.5 embryos and 21-day-old pups of each genotype obtained by crossing heterozygous *Sned1^WT/-^* mice (\**1 mouse died atp22 and another one at p24*).

| *Sned1*<sup>WT/-</sup> x *Sned1*<sup>WT/-</sup> | *Sned1*<sup>WT/WT</sup> | *Sned1*<sup>WT/-</sup> | *Sned1*<sup>-/-</sup> | $\chi^2$<br>p-value |
| --- | --- | --- | --- | --- |
| **Expected** | 25% | 50% | 25% |  |
| **E18.5** | 12 (27.3%) | 20 (45.5%) | 12 (27.3%) | 0.834 |
| **Alive at weaning (p21)** | 14 (22.2%) | 44 (69.8%) | **5\* (7.9%)** | **0.0019** |

5 litters were harvested by C-section to examine a total of 44 E18.5 embryos. As observed for the *Sned1^LacZ-Neo^* strain, Mendelian ratios were observed (χ^2^=0.364; two-tailed p=0.834) indicating that lethality of *Sned1*^-/-^ pups occurred around or shortly after birth (Table 2).

Altogether, our data demonstrate that *Sned1* knockout, achieved using two approaches, results in early neonatal lethality, with around 67% penetrance (Tables 1 and 2).

Thus, over the course of our study, some knockout mice survived (Tables 1 and 2). Remarkably, the knockout survivors (either *Sned1^LacZ-Neo/LacZ-Neo^* or *Sned1*^-/-^) were easily distinguishable from wild-type and heterozygous littermates because of their smaller size and the shape of their head and snout. We thus sought to characterize in detail their phenotype to gain insight into the physiological roles of SNED1. Of note, crossing of *Sned1^LacZ-Neo/LacZ-Neo^* survivors revealed that surviving *Sned1* knockout mice were fertile although they produced less numerous and smaller litters. However, pups obtained by breeding *Sned1^LacZ-Neo/LacZ-Neo^* survivors displayed exacerbated phenotypes (in particular, even shorter snouts, under-developed or lacking mandibles; not shown) and only some of their progeny survived after weaning.

### *Sned1* knockout results in growth defects

Our first observation was that knockout survivors (either *Sned1^LacZ-Neo/LacZ-Neo^* or *Sned1*^-/-^) were visibly smaller than wild-type and heterozygous littermates (Figure 3A). At p14, knockout pups weighed significantly less than wild-type and heterozygous littermates (Figures 3B and 3C). This reduced body weight was readily detectable in p0.5 neonates (data not shown). Suspecting skeletal anomalies, we acquired micro-computed tomography (μCT) scans of these mice when they reached adulthood and measured the length of their long bones.

**Figure 3.**
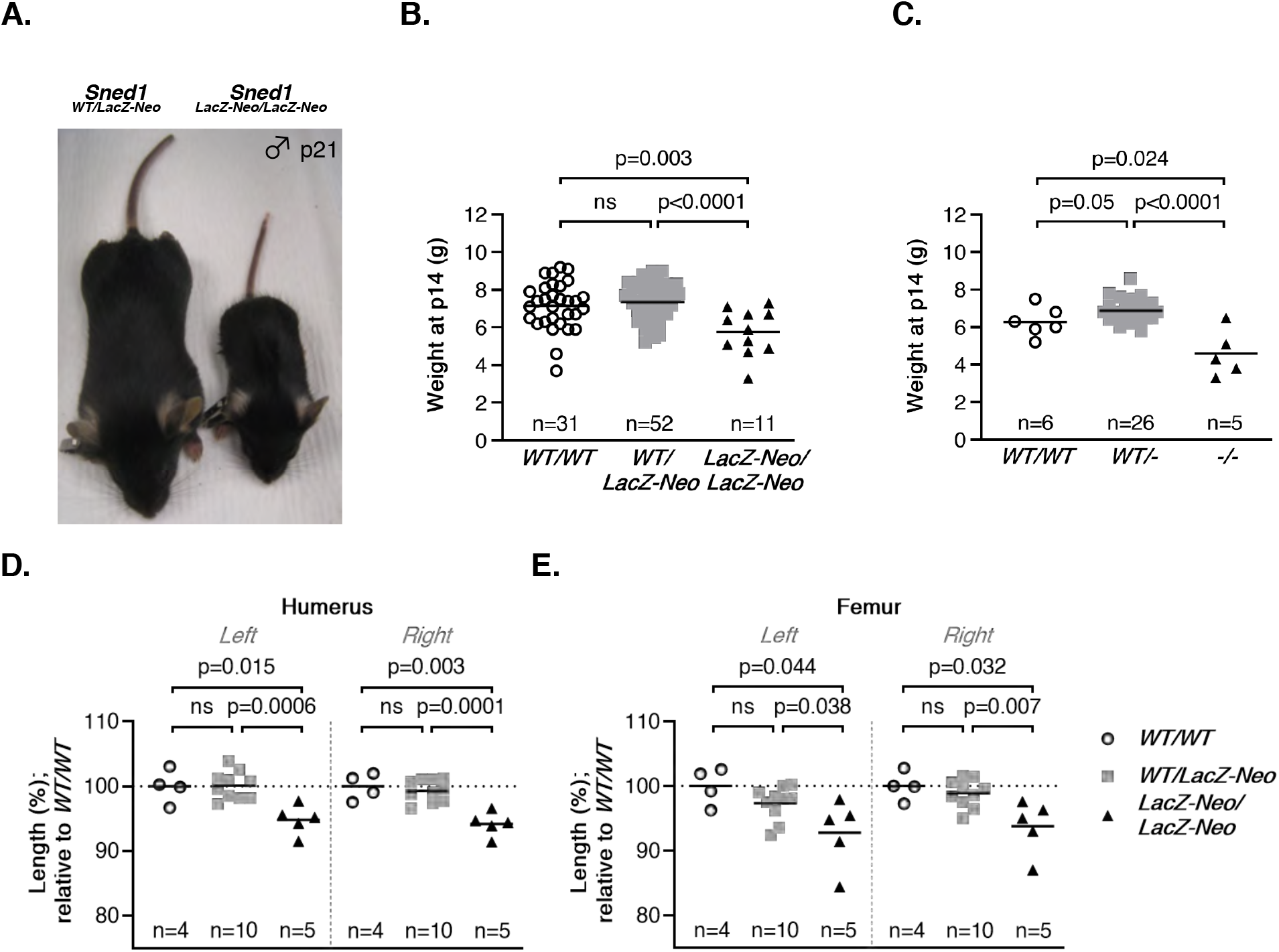
Surviving *Sned1* knockout mice present growth defects. **A.** Representative pictures of heterozygous (left) and knockout (right) 21-day-old mice illustrate the smaller size of surviving knockout mice. **B-C.** Dotplots represent the weight of 14-day-old pups of the *LacZ-Neo* transgenic line (**B**) or null transgenic line (**C**). Numbers of mice per genotype are indicated. Line represents average weight of the genotype group. Statistical analysis was performed using an unpaired t-test. **D-E.** Dotplots represent the length of the humerus (D) and femur (E) of adult mice of the *LacZ-Neo* transgenic line. Numbers of mice per genotype are indicated. Line represents average weight of the genotype group. Statistical analysis was performed using an unpaired t-test.

We found that *Sned1^LacZ-Neo/LacZ-Neo^* showed statistically significant bilateral shortening of the humerus and femur compared to control and heterozygous littermates (Figures 3D and 3E) and of all other bones measured (ulna, radius, and tibia; data not shown). No significant differences were found between the wild-type and heterozygous groups.

### *Sned1* knockout mice present craniofacial malformations

Besides their small size, knockout survivors were also distinguishable from wild-type and heterozygous littermates by their shorter snouts and abnormal head shape. Thus, we examined the skulls of wild-type, heterozygous and *Sned1* knockout mice using μCT (Figure 4A). We measured the length of the different calvarial sutures including the nasal, frontal, and sagittal sutures (Figure 4A). Knockout mice showed statistically significantly shorter nasal sutures as compared to wild-type (p=0.004) or heterozygous (p=0.005) mice (Figure 4B, left panel). Frontal sutures did not differ significantly in length, although there was a trend towards shorter sutures in the knockout mice (Figure 4B, right panel). The length of the sagittal sutures, joining parietal bones of mesodermal origin, although tending to be shorter, did not show statistically significant differences between groups (data not shown).

**Figure 4.**
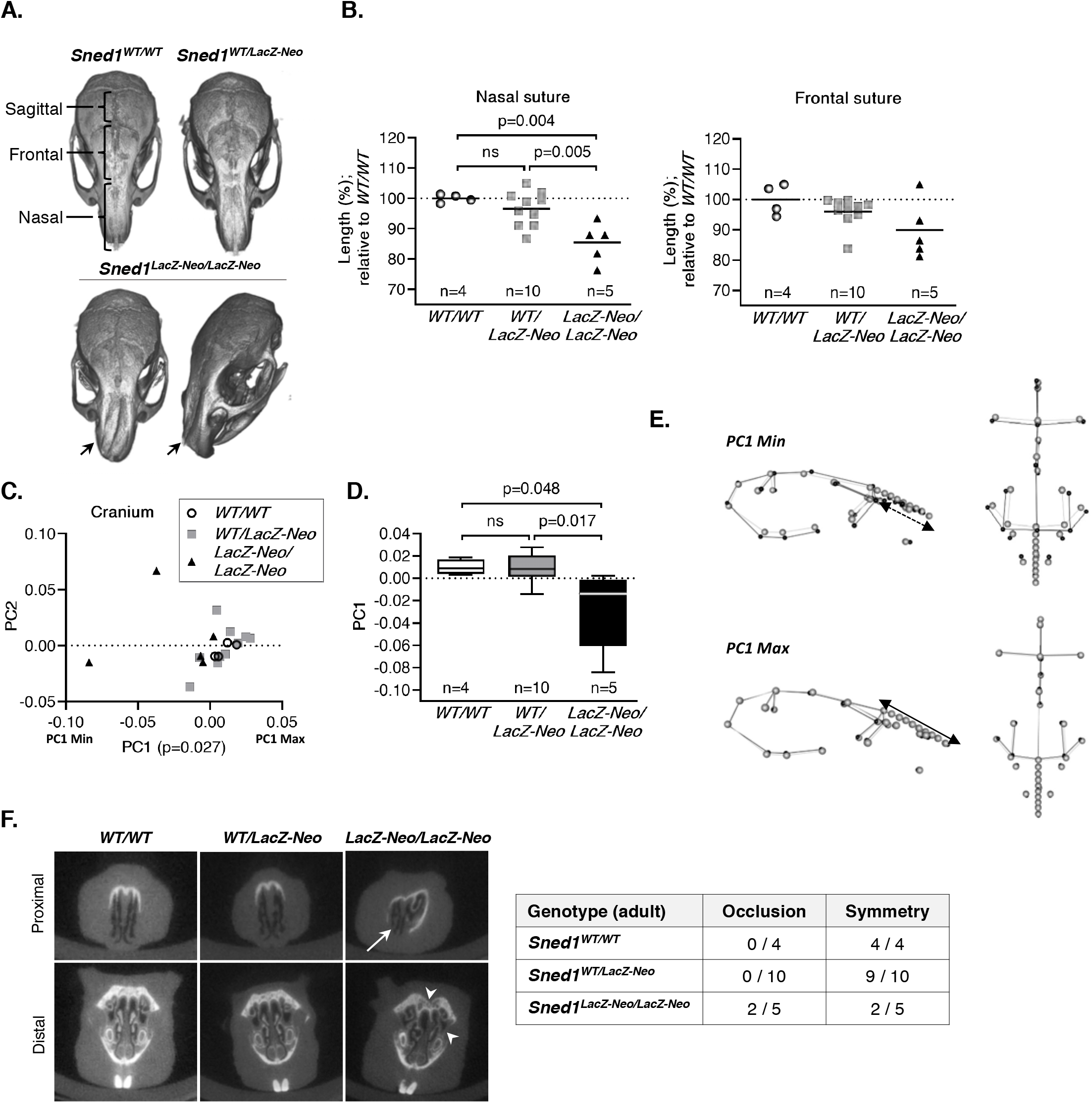
*Sned1* knockout results in craniofacial malformations. **A.** μCT scans showing variation in skull morphology based upon genotype. Note the nasal bridge collapse in *Sned1^LacZ-Neo/LacZ-Neo^* mouse (arrow). **B.** Nasal (left panel) and frontal (right panel) suture length. Numbers of mice per genotype are indicated. Line represents average length of the genotype group. Statistical analysis was performed using an unpaired t-test. **C.** Scatterplot of PC1 and PC2 for variation in the mouse cranial morphology. Statistical analysis was performed using a one-way ANOVA. **D.** Boxplot of cranial PC1 data by mouse genotype. Numbers of mice per genotype are indicated. Statistical analysis was performed using a Tukey’s HSD test. **E.** Wireframes depicting mouse cranium side (left) and top (right) views and variation of the shape as a function of genotypes. Grey dots represent the consensus cranium shape, black dots represent the shape depicted by maximum and minimum PC1 range. **F.** Occlusion and asymmetry assessment of the nasal cavity of 19 adult (6-month-old) mice. Arrow indicates occlusion and arrowheads indicate asymmetries.

We further employed geometric morphometric analysis, an approach that compares the relative positions of anatomical landmarks^41^, to quantify complex 3-dimensional shape variations in between mice of different genotypes. To do so, we positioned 22 traditional coordinate landmarks which were deemed to represent well overall cranial morphology and an additional 8 sliding semi-landmarks along the nasal suture, since this structure was particularly affected upon *Sned1* knockout (see Supplementary Method 1). Landmark configurations are then superimposed and compared using principal component analysis (PCA). PC A yielded 19 principal components, the first four of which represented over 5% of the overall variation. The first principal component (PC1) accounted for 30.05% of the overall variation and described differences in cranial vault shape and nasal bridge morphology. PC1 showed statistically significant differences across the three groups (p=0.027, Figure 4C). A Tukey’s honest significant difference (HSD) test revealed that the overall statistical significance in PC1 was driven by differences between the wild-type and knockout groups (p=0.048) and between the heterozygote and knockout groups (p=0.017) while the comparison of the heterozygous and wild-type mice was not statistically significantly different (p=0.99) (Figure 4D). Wireframes depicting the consensus cranium shape was drawn (Figure 4E, grey dots) and the average shapes of the extreme ends of the range of variation for PC1 were superimposed (Figure 4E, black dots). Animals at the minimum extreme of variation along PC1 (PC1 min, knockout mice) showed concavity of the nasal bridge, relatively shorter snouts, wider faces and neurocrania, and more supero-inferiorly expanded neurocrania (Figure 4E, upper panel).

In contrast, those at the maximum extreme of variation along PC1 (PC1 max, wild-type and heterozygous mice) tended to show relatively flatter (supero-inferiorly) and medio-laterally narrower neurocrania, narrower midfaces (at the zygoma), and a more convex nasal bridge (Figure 4E, lower panel). Although the mechanisms by which SNED1 controls nasal bridge formation remain unknown, we can hypothesize that they contribute to the proper formation of the nasal septum, the cartilaginous structure that supports the ossified nasal bridge.

### *Sned1* knockout mice present asymmetric and occluded nasal cavities

The observation that *Sned1* knockout mice present nasal bridge collapse prompted us to further evaluate the anatomy of the nasal cavity of the animals. In the cohort of adult mice, 2 out of the 5 knockout mice presented partial occlusion, defined as a contact between the soft tissues forming the wall of the airways, whereas none of the wild-type and heterozygous mice presented occlusion. In addition, 3 out of the 5 knockout mice presented markedly asymmetric nasal cavities, whereas 9 out of 10 heterozygous and all 4 wild-type mice had symmetrical cavities (Figure 4F, Supplementary Files 1-3). Interestingly, nasal occlusion is already present at birth, since the evaluation of a small cohort of neonates (p0.5) revealed that all knockout neonates analyzed presented at least partial nasal occlusion, in contrast to none of the wild-type animals, and only 3 out of the 15 heterozygotes studied (Supplementary Files 4-6). Asymmetric nasal cavities and airway obstruction support the hypothesis that the *Sned1* knockout neonates may not survive because of impaired nasal respiration.

### *Sned1* knockout mice present under-developed mandibles

Having observed occasionally that some of the *Sned1*^-/-^ neonates, who did not survive, presented severely affected jaws (Figure 5A), and having observed that *Sned1* is expressed in Meckel’s cartilage from which the mandibles develop (Figure 6D, *see below*), we further investigated variations in the mandibular structure using geometric morphometric analysis. PCA yielded 18 principal components, the first three of which represented greater than 5% of the overall variation. PC1 represented 26.51% of the variation and showed statistically significant differences across the three groups (p<0.0001) (Figure 5B). A Tukey’s HSD test revealed that these differences were statistically significant between the wild-type and knockout groups (p<0.0001) and between the heterozygote and knockout groups (p<0.0001) while the comparison of the heterozygous and wild-type mice was not statistically significantly different (p=0.679) (Figure 5C).

**Figure 5.**
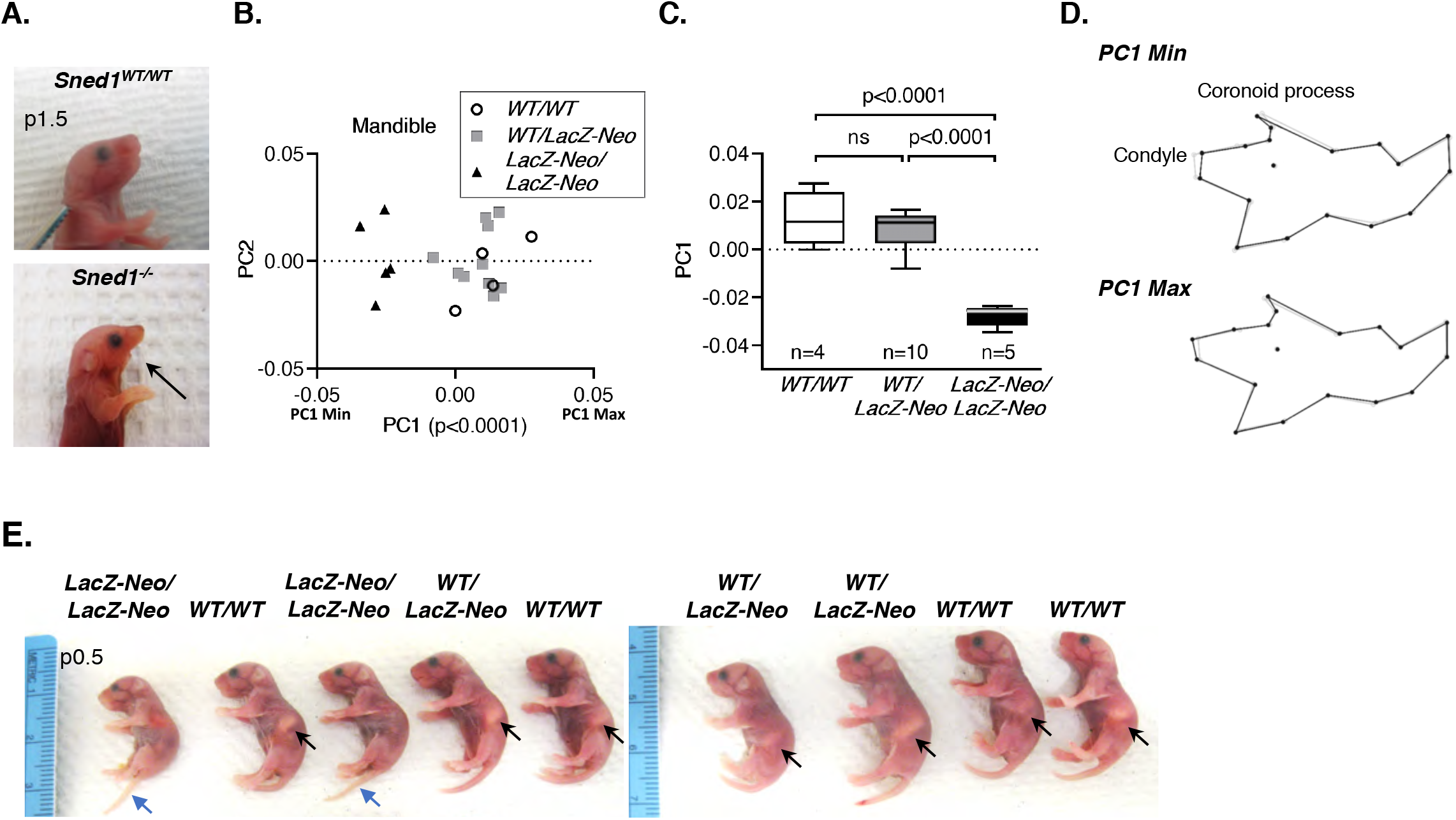
*Sned1* knockout results in under-developed mandibles. **A.** Pictures of a p1.5 wild-type (upper panel) and age-matched dead *Sned1^-/-^* (lower panel) neonates showing the absence of the lower jaw in the knockout pup. **B.** Scatterplot of PC1 and PC2 for variation in the mouse mandible morphology. **C.** Boxplot of mandibular PC1 data by mouse genotype. Numbers of mice per genotype are indicated. Statistical analysis was performed using a Tukey’s HSD test. **D.** Wireframes depicting mouse mandible side view and variations as a function of genotypes. Grey dots represent the consensus cranium shape, and black dots represent the shape depicted by minimum (upper panel) and maximum (lower panel) PC1 range. **E.** Representative litter of p0.5 neonates. Black arrows indicate the presence of milk the stomach of wild-type and heterozygous mice. Note the absence of milk in the stomach of knockout neonates. Blue arrow points to the tails of knock-out neonates which fail to curl, perhaps related to the strong expression in tail-bud somites and mesoderm (*Fig 6*).

**Figure 6.**
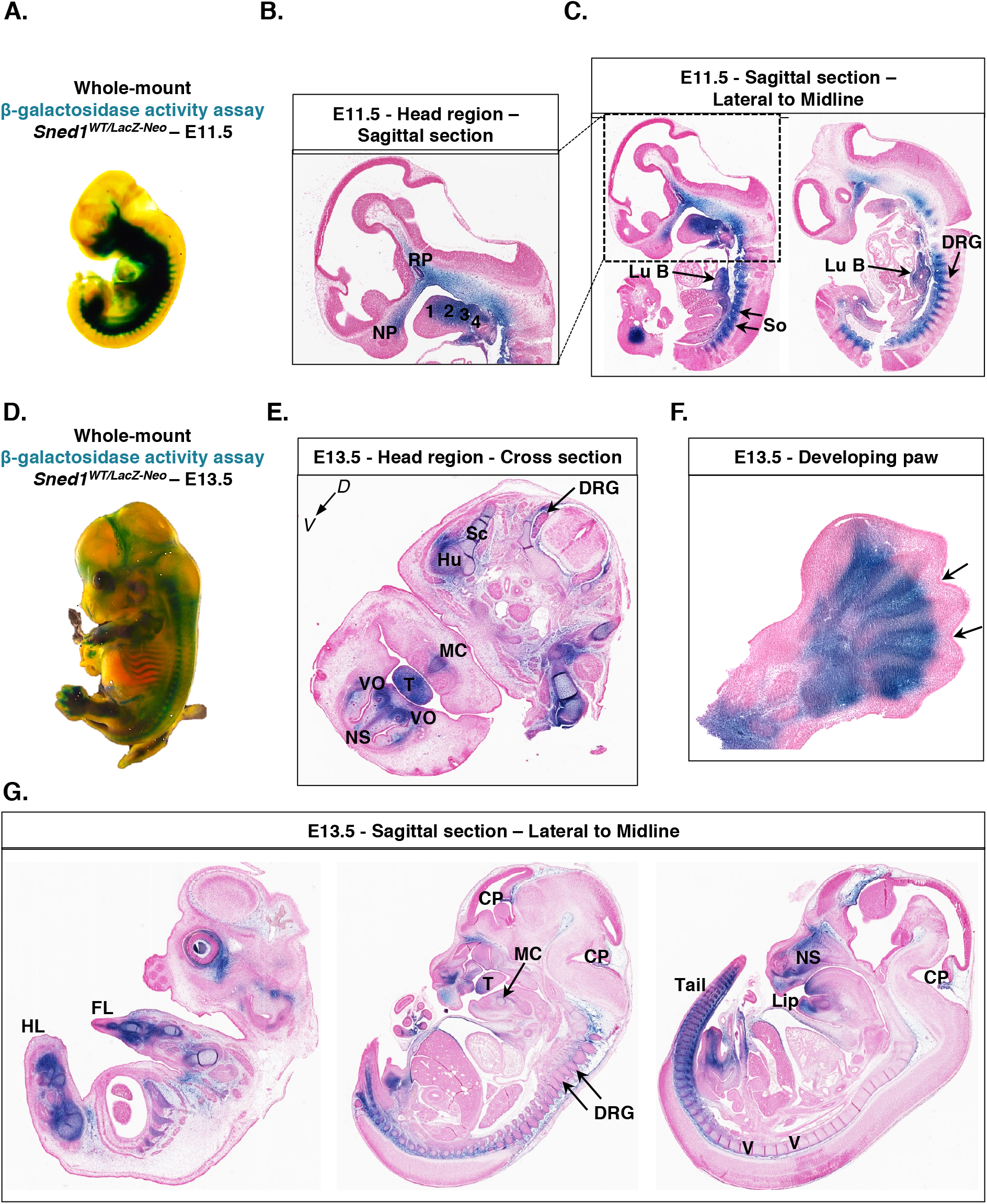
Patterns of expression of *Sned1* gene during embryogenesis. **A.** Whole-mount β-galactosidase assay (LacZ staining) performed on heterozygous E11.5 *Sned1^WT/LacZ-Neo^*embryo. **B.** Sagittal section of the head region of LacZ-stained E11.5 LacZ *Sned1^WT/LacZ-Neo^* embryo. 1, 2, 3, and 4 represent the branchial arches; NP: nasal process; RP: Rathke’s pouch. **C.** Sagittal sections of LacZ-stained E11.5 LacZ *Sned1^WT/LacZ-Neo^*embryo (left panel: lateral section; right panel: across the midline). Lu B, lung bud; So, somites; DRG, dorsal root ganglia. **D.** Whole-mount β-galactosidase assay (LacZ staining) performed on heterozygous E13.5 *Sned1^WT/LacZ-Neo^*embryo. **E.** Transverse section of the head region of LacZ-stained E13.5 LacZ *Sned1^WT/LacZ-Neo^* embryo. DRG, dorsal root ganglia; Hu: humerus; MC: site of apposition in the midline of Meckel’s cartilages; NS, cartilage primordium of nasal septum; Sc: blade of scapula; T, tongue; VO, vomeronasal organ. **F.** Section of paw from LacZ-stained E13.5 LacZ *Sned1^WT/LacZ-Neo^* embryo. Arrows indicate interdigital spaces. **G.** Sagittal sections of Lac-Z-stained E13.5 LacZ *Sned1^WT/LacZ-Neo^* embryo (from left to right: lateral to the midline). CP, choroid plexus; DRG, dorsal root ganglia; FL, forelimb; HL, hindlimb; NS, cartilage primordium of nasal septum; T, tongue; V, cartilage primordium of the vertebrae.

Mice at the minimum extreme of variation along PC1 (PC1 min, knockout mice), show shorter mandibular condyles and smaller coronoid processes (Figure 5D, upper panel). In contrast, those at the maximum extreme of variation along PC1 (PC1 max, wild-type and heterozygous mice) tend to display longer mandibular condyles and larger coronoid processes (Figure 5D, lower panel). Interestingly, we also observed that no milk was present in the stomachs of p0.5 knockout neonates whereas milk was present in wild-type and heterozygous littermates (Figure 5E). Although the mechanisms by which SNED1 controls mandible formation remain unknown, we hypothesize that they may affect knockout mice’s ability to suckle or chew properly, as the mandibular condyle and coronoid process are attachment points for chewing muscles such as the masseter. The mandibular condyle is also an important growth site of the mandible, contributing to the elongation of the ramus.^42^. It will thus be interesting to determine whether the growth defect of *Sned1* knockout mice (Figure 3) is an indirect result of the decreased ability of knockout mice to eat or whether SNED1 plays a direct role during skeletal development.

### *Sned1* is expressed in skeletal and craniofacial precursors

We took advantage of the LacZ reporter gene expressed under the control of the *Sned1* promoter to determine in which tissues *Sned1* was expressed during development. Whole-mount LacZ staining revealed that the *Sned1* promoter is broadly active at the two time-points examined, embryonic days 11.5 (Figure 6A) and 13.5 (Figure 6D). Many of the zones of expression conform with tissues that later show defects. At E11.5, LacZ staining was observed in the nasal process and the pharyngeal arches (Figure 6B), structures populated by cephalic neural crest cells (NCCs) that will contribute to form the skull and facial skeleton.^43,44^ At E11.5, LacZ staining was also detected in mesodermal derivatives including the ventro-medial cells of the somites (Figure 6C), which will also undergo epithelial-to-mesenchymal transition (EMT) to form the sclerotomes which will, in turn, give rise to the vertebrae and the ribs. In addition, staining was detected in the lung bud (Figure 6C), as previously reported^22^ and in Rathke’s pouch, a structure deriving from the oral ectoderm which will give rise to the adenohypophysis of the pituitary gland. At E13.5, staining was visible in the head region, in the derivatives of the pharyngeal arches, including the tongue and the Meckel’s cartilage, which will contribute to the development of the mandible, the cartilage primordium of the nasal septum, and the vomeronasal organ, a chemoreceptive organ part of the olfactory system (Figures 6E and 6G). Staining was also observed in the skeletal precursors of the humeral shaft and the scapula (Figure 6E), the vertebrae and the ribs (Figures 6D and 6G). Of note, the LacZ staining intensity followed the anteroposterior axis, with the posterior somites being the last to differentiate and the most intensely stained (Figure 6G, right panel). Staining was also observed in the limb buds (Figures 6F and 6G), the choroid plexus (Figure 6G, middle and left panel), and in the mesenchyme surrounding the dorsal root ganglia. Our results confirm and extend the previously published study reporting the expression of *Sned1* using *in-situ* hybridization,^22,45^ and with gene expression data reporting *Sned1* expression in the mandible and frontonasal region of E10.5 to E12.5 embryos.^46,47^

### Deletion of *Sned1* in the neural crest cells does not affect mouse survival

The craniofacial skeleton has two embryonic origins; the cephalic NCCs form the anterior region of the skull, while the mesoderm contributes to the more posterior cranial bones.^48,49–47,48^ Interestingly, the length of the sagittal suture, formed by the parietal bones of mesodermal origin, was not statistically different between wild-type, heterozygous, and *Sned1* knockout mice (data not shown). Following induction, cephalic NCCs undergo EMT and migrate from the dorsal neural tube into the frontonasal region and pharyngeal arches, giving rise to most of the bone and cartilage of the head and neck.^25–28^ Of note, the mesenchyme of nasal, frontal, and sagittal sutures is also neural-crest derived.^50^ Because *Sned1* is expressed in structures that will form the craniofacial skeleton and NCC-derived tissues, including the nasal bone and the mandible, and because these structures are abnormal in *Sned1* knockout mice, we hypothesized that it may regulate NCC phenotype. Interestingly, the origin of vertebrates is concomitant with the appearance of the NCC population,^36,51^ and orthology analysis has found that orthologues of *Sned1* are found in all sequenced vertebrates, but not in lower organisms (Figure 1).

To assess the putative role of SNED1 in the NCCs, we ablated *Sned1* specifically from NCCs. To do so, we crossed mice carrying the conditional *Sned1* allele (*Sned1^fl/fl^*) with the previously characterized and widely used *Wnt1-Cre* transgenic mouse line^52^ (Figure 7A). We examined 32 litters born from 6 different breeding pairs (representing a total of 180 pups). Contrary to what we observed in the global *Sned1* knockout mice, the NCC-specific deletion of *Sned1* did not affect survival since mice were present in Mendelian ratios (χ^2^=5.378; two-tailed p=0.146) at weaning (p21 +/- 1 day, Table 3). These data suggest that the production of SNED1 by other cell types apart from the NCCs is essential for development.

**Figure 7.**
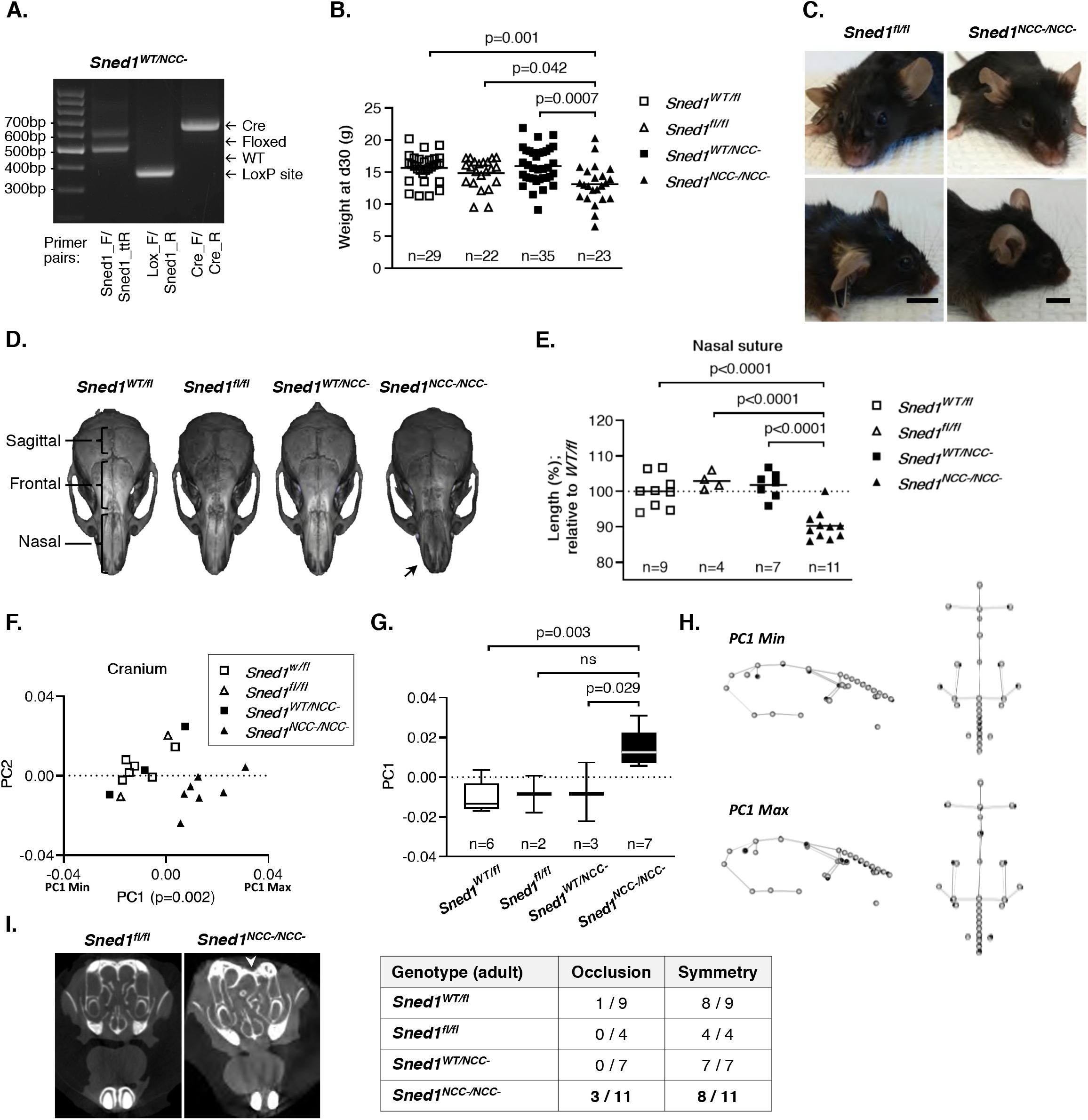
NCC-specific *Sned1* knockout results in craniofacial malformations. **A.** PCR genotyping of genomic DNA isolated from tail samples of *Sned1^WT/NCC^* mice. **B.** Dotplots represent the weight of one-month-old mice of the *Sned1^NCC^* transgenic line. Numbers of mice per genotype are indicated. Lines represent the average weight of the genotype group. Statistical analysis was performed using an unpaired t-test. **C.** Representative pictures of 9-month-old control (*Sned1^fl/fl^*, left panels) and knockout (*Sned1^NCC-/NCC^*, right panels) mice. **D.** μCT scans show examples of variation in skull morphology based upon genotype. **E.** Nasal suture length. Numbers of mice per genotype are indicated. Lines represent average length of the genotype group. Statistical analysis was performed using an unpaired t-test. **F.** Scatterplot of PC1 and PC2 for variation in the mouse cranial morphology of the *Sned1^NCC-^* transgenic line. Statistical analysis was performed using a one-way ANOVA. **G.** Boxplot of cranial PC1 data by mouse genotype. Numbers of mice per genotype are indicated. Statistical analysis was performed using a Tukey’s HSD test. **H.** Wireframes depicting mouse cranium side (left) and top (right) views and variation of the shape as a function of genotypes. Grey dots represent the consensus cranium shape, black dots represent the shape depicted by maximum and minimum PC1 range. **I.** Occlusion and asymmetry assessment of the nasal cavities of 31 adult (7 to 15-month-old) mice. Arrowhead indicates asymmetry.

**Table 3.** Numbers of expected and observed 21-day-old pups of each genotype obtained by crossing *Wnt1-Cre; Sned1^fl/fl^* with *Sned1^fl/fl^* mice.

| *Wnt1-Cre; Sned1<sup>WT/fl</sup>* x *Sned1<sup>fl/fl</sup>* | *Sned1<sup>WT/fl</sup>* | *Sned1<sup>fl/fl</sup>* | *Wnt1-Cre; Sned1<sup>WT/fl</sup>*<br>(*Sned1<sup>WT/NCC-</sup>*) | *Wnt1-Cre; Sned1<sup>fl/fl</sup>*<br>(*Sned1<sup>NCC-/NCC-</sup>*) | $\chi^2$<br>*p-value* |
| --- | --- | --- | --- | --- | --- |
| **Expected** | 25% | 25% | 25% | 25% |  |
| **Alive at weaning (p21)** | 57 (31.7%) | 36 (20.0%) | 46 (25.5%) | 41 (22.8%) | 0.146 |

### Deletion of *Sned1* in the neural crest cells results in growth defects and craniofacial malformations partly phenocopying the effect of the global deletion of *Sned1*

Since the global *Sned1* knockout resulted in growth defects, we also recorded possible differences in body weight and observed that one-month-old *Sned1^NCC-/NCC-^* mice were smaller and weighed less than control and heterozygous littermates (Figure 7B). Importantly, and similarly to global knockout mice, *Sned1^NCC-/NCC-^* mice could be easily distinguished because they presented craniofacial malformations, including shorter snouts and wider neurocranium (Figure 7C). To further examine variations in shape based on the genotype, we scanned the skulls of adult mice using μCT. We observed that the nasal bone was severely malformed upon deletion of *Sned1* in the NCCs (Figure 7D, arrow). *Sned1^NCC-/NCC-^* mice showed significantly shorter nasal sutures as compared to controls (*Sned1^WT/fl^* and *Sned1^fl/fl^*) and heterozygotes (*Sned1^WT/NCC^*) (Figure 7E). No statistically significant difference was observed in the length of the frontal suture between genotype groups, although there was a trend toward *Sned1^NCC-/NCC-^* mice having shorter frontal sutures. Contrary to what we observed upon global *Sned1* knockout, sagittal sutures of *Sned1^NCC-/NCC-^* mice were also shorter as compared to control and heterozygous mice (data not shown).

We then applied the same standard geometric morphometric analysis as the one employed to characterize the craniofacial features of the global KO mice, to assess variations in the shape of the skulls of a subset of our cohort of control and NCC-specific knockout mice (6 *Sned1^WT/fl^*, 2 *Sned1^fl/fl^*, 3 *Sned1^WT/NCC-^* and 7 *Sned1^NCC-/NCC-^*). Principal component analysis yielded 17 PCs, the first 5 of which represented at least 5% of the total variation each. Analysis of the variance of the 4 different groups along the first 5 PC axes found statistically significant differences along PC1 (p=0.002; Figure 7F). A Tukey HSD test further revealed that *Sned1^NCC-/NCC-^* mice drove this variation, and was statistically significantly different from control (p=0.003) and heterozygote mice (p=0.029). As shown Figure 7F, *Sned1^NCC-/NCC-^* mice cluster at the positive end of the range of variation along PC1 (PC1 max), which represents mice with more concave nasal bridges, relatively shorter snouts, and relatively wider crania (see wireframes Figure 7H); the same morphological craniofacial features observed upon global *Sned1* deletion.

Last, we evaluated the anatomy of the nasal cavity in the NCC-specific *Sned1* knockout mice. Out of the 11 knockout mice, 3 presented partial occlusion of the nasal cavities. In contrast, only one of the controls (*Sned1^WT/fl^*) and none of the heterozygous mice showed airway occlusion. We also observed that 3 knockout mice presented severely asymmetric nasal cavities whereas 12 out of the 13 controls (8 *Sned1^WT/fl^* and 4 *Sned1^fl/fl^*) and all 7 heterozygotes had symmetrical cavities (Figure 7I, Supplementary Files 7-11).

Altogether, our results demonstrate that the deletion of *Sned1* in the NCCs partly phenocopies what we observed in global knockout mice, providing evidence that SNED1 is important for the NCC population and, eventually, proper craniofacial morphogenesis. The fact that *Sned1^NCC-NCC-^* mice survive could be explained by the fact that SNED1 might be acting in a cell-autonomous manner on mesodermal-derived structures contributing to the formation of the facial skeleton. In addition, SNED1 produced by other cell types might function as signaling cues for NCCs, ultimately ameliorating the severity of the phenotype in *Sned1^NCC-NCC-^* mice.

## Discussion

We report here the generation and phenotypic analysis of mouse strains in which we deleted *Sned1* globally or specifically from the neural crest cells. We showed that *Sned1* is essential during development, since the majority of the global knockout mice died prematurely within 2 days of birth, and the surviving animals exhibit severe growth defects and craniofacial malformations. Despite the broad pattern of expression of *Sned1* in murine embryos, the only obvious phenotypes discerned at this stage of our studies relate to overall growth, long bone length, and craniofacial features. The surviving *Sned1* knockout adults did not present other overt anomalies (for example, normal number of digits was counted, tongue appeared normal, complete palate closure was observed, no pigmentation defect was observed). It is worth noting that the adult mice on which the phenotypic analyses were conducted are likely to present less severe phenotypes because they survived. Follow-up studies will be required to understand whether this is stochastic or whether compensatory mechanisms have intervened and led to certain animals surviving whereas the majority of *Sned1* knockout neonates did not survive past post-natal day 2.

Based on *Sned1* expression data and the observed craniofacial malformations, we postulated that SNED1 may play a role in NCC biology. Interestingly, deletion of *Sned1* in the NCCs also results in growth defects and facial dysmorphic features, partly phenocopying the global knockout. We previously identified the function of SNED1 as an ECM protein promoting breast cancer metastasis.^5^ Here, we report the expression of *Sned1* in structures that undergo EMT and cell migration during embryonic development, including the sclerotomes and the NCCs.^53^ The parallel has been drawn between developmental EMT and carcinoma metastasis, since in both cases, epithelial cells lose cell-cell junctions, up-regulate ECM genes such as fibronectin, remodel their adhesions to the ECM and acquire a migratory phenotype.^54,55^ Therefore, the NCC-specific *Sned1* KO will be a useful tool to decipher which precise cellular mechanism(s) governing NCC behavior (EMT, migration, proliferation, differentiation) are controlled by SNED1. Studies of the possible roles of SNED1 in EMT of NCC may thus also further help understand how SNED1 contributes to cancer metastasis.

Once cephalic NCCs have migrated into the frontonasal region and pharyngeal arches, they need to receive signals in order to survive, proliferate, and differentiate into bone and cartilage. Future studies will also evaluate whether SNED1 contributes to any of these processes.

As of now, neither the SNED1 receptor(s) nor the signaling pathways activated downstream of SNED1 are known. Dissecting the molecular mechanisms leading to the phenotypes described in the present study will be the focus of future investigations. In particular, integrins have been shown to be involved in different aspects of cranial neural crest cells phenotypes. ^17,56–61^ It would thus be important to determine whether the putative integrin-binding sites found in SNED1 do indeed engage these ECM receptors, and if so, whether this is how SNED1 governs craniofacial morphogenesis. In addition, it will be crucial to identify the pathways regulating SNED1.

Of note, *Sned1* was among the most significantly up-regulated genes in NCC-specific *Tgfbr2* knockout mice (*Wnt1-Cre; Tgfbr2^fl/fl^*) presenting abnormal palate closure (clefting)^62^ (Table 4). Although cleft palate or lip were not observed in *Sned1* knockout mice, this observation may suggest that SNED1 is regulated by the TGF-β signaling pathways. Supporting this idea, *Sned1* was found to be enriched upon removal of *Tgfbr2* in the developing intervertebral discs (*Co12a1-Cre; Tgfbr2^fl/fl^*) of mouse embryos, and down-regulated when sclerotome cells are treated with ligands of the TGF-β superfamily^63^ (Table 4). The TGF-β signaling pathway plays fundamental roles in bone and cartilage formation and homeostasis,^64^ but also in metastasis.^65^ A literature search also revealed that *Sned1* is enriched in the mandible and frontonasal region of mouse embryos,^46,47^ murine differentiating osteoblasts,^66^ and human articular chondrocytes of fetal joints^67^ (Table 4), suggesting a putative role for SNED1 in chondrogenesis and osteogenesis in bones of the axial and appendicular skeleton. Future mechanistic investigations will benefit from the conditional *Sned1* knockout mouse generated in this study which will be instrumental in determining the functions of *Sned1* in a cell- or tissue-specific and time-specific manner.

**Table 4.**
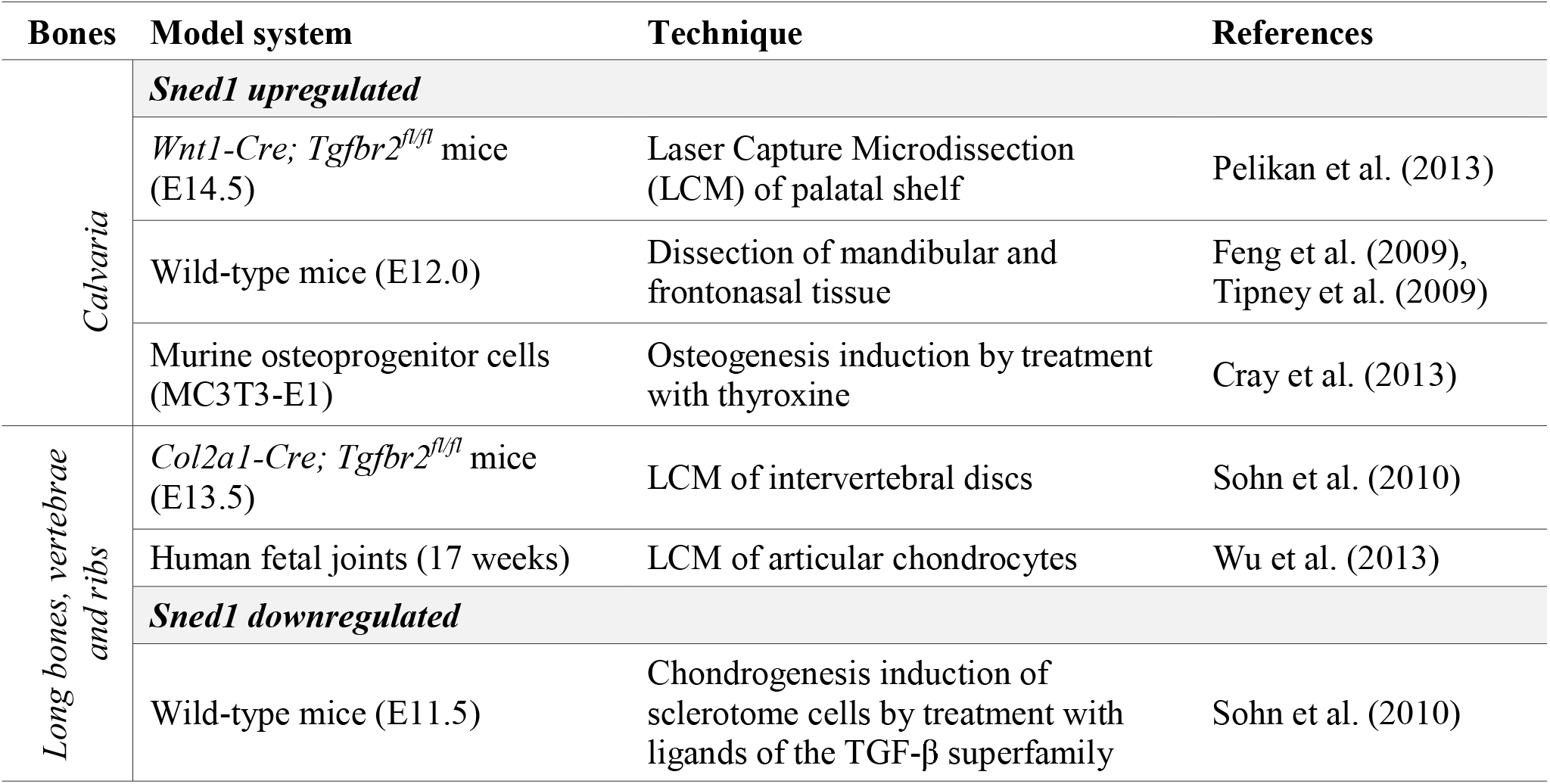
Studies reporting *Sned1* in high-throughput microarray datasets of skeletal tissues

Last, future studies of the roles of SNED1 in craniofacial development using the mouse models generated in this study may shed light on fundamental processes that, when altered, lead to craniofacial malformations in humans.^68^ *SNED1* is localized on the long arm of chromosome 2 (2q37.3) in humans. Patients with 2q37 deletion syndrome, also described in the literature as Albright hereditary osteodystrophy-like syndrome or brachydactyly-mental retardation (BDMR) syndrome, suffer from mental retardation, facial dysmorphism, including round face and flattened nasal bridge, and skeletal abnormalities such as brachydactyly and short stature.^69–71^ More than 100 cases have been reported in the literature with 2q37 deletion syndrome.^72^ Genotype-phenotype correlation studies have been performed to delineate the critical region causing BDMR.^72^ *HDAC4* has been postulated as one of the candidates responsible for BDMR.^73^ However, at least two cases with facial dysmorphic features have been reported harboring a distal 2q37.3 deletion not including *HDAC4*^73,74^ (Figure 8). Since *SNED1* had not been identified when most of these studies were published and that the chromosomal breakpoints had not been precisely mapped in patients diagnosed more than 20 years ago,^75,76^ there are potentially more patients presenting with no deletion of *HDAC4* but with deletion of *SNED1*. In light of the present study, it would thus be interesting to determine to what extent the craniofacial phenotype of 2q37.3 deletion syndrome patients can be attributed to the loss of *SNED1*.

**Figure 8.**
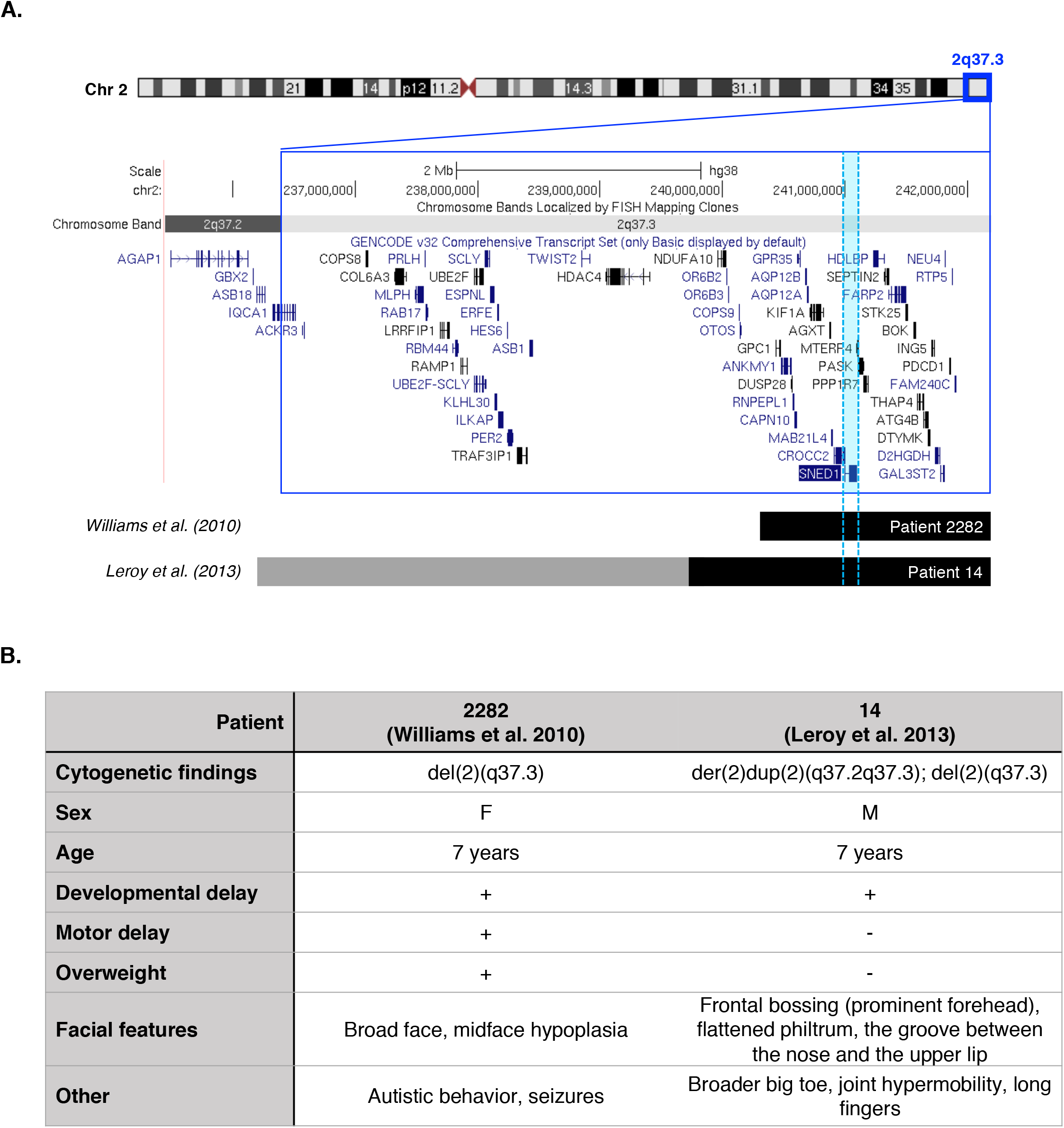
Map of 2q37.3 deletions in two patients with facial dysmorphic features. **A.** Schematic representation of the human 2q37 chromosome region (dark blue box) showing two patients carrying a terminal deletion that includes the *SNED1* locus (highlighted in light blue) but not *HDAC4*. Deleted regions are indicated by horizontal black bars and gray bar indicates duplicated region in patient 14 from the Leroy at al, 2013 study. *Source:* UCSC genome browser: http://genome.ucsc.edu, GRCh38/hg38 Assembly. **B.** Clinical features of two patients with 2q37.3 deletion including *SNED1* but not *HDAC4*.

## Experimental Procedures

### Generation of *Sned1* knockout mice

All experiments involving mice were conducted in conformity with animal protocols approved by MIT’s Department of Comparative Medicine and Committee on Animal Care, and by the Committee on Animal Care of the University of Illinois at Chicago in compliance with IACUC guidelines.

Two targeted *Sned1^tm1a(KOMF)Wtsi^* ES cell clones (EPD0300_5_02 E02 and F01) from the C57BL/6 JM8.N4 parental ES cell line generated by the Wellcome Trust Sanger Institute^40^ were obtained from the KOMP Repository (www.komp.org) at UC Davis and cultured on feeder cells using the recommended media including the 2i additives MEK inhibitor PD0325901 and GSK3 inhibitor CHIR99021 (StemGent, Cambridge, MA). Three subclones of each clone were sent to Cell Line Genetics (Madison, WI) for cytogenetic analysis and two subclones of the targeted clone E02 were found to have an apparently normal mouse male karyotype. Cells from subclone E02-2D2 were microinjected into BALB/c blastocysts obtained from natural matings as previously described.^77^ 10 recipient females each received 14 embryos. 7 pregnant females gave birth to 41 pups, of which 6 showed at least 70% dark coat color against white, BALB/c fur, and allowed establishment of the *Sned1^tm1a(KOMP)Wtsi^* (referred to as *Sned1^LacZ-Neo^*) mouse line. B6.Cg-Tg(ACTFLPe)9205Dym/J (Actin-Flp; Jackson Laboratory stock No. 005703) mice allowing the ubiquitous expression of the Flp recombinase were obtained from MIT’s transgenic core facility and were used to generate the *Sned1* floxed *Sned1^mlc(KOMP)WtsI^* allele, referred to as the *Sned1^fl^* allele. B6.C-Tg(CMV-cre)1Cgn/J (CMV-Cre) mice allowing the ubiquitous expression of the Cre recombinase were purchased from Jackson Laboratory, stock No. 006054 and were used to generate the null *Sned1^tm1d(KOMP)Wtsi^* allele, referred to as the *Sned1* allele. Figure 2A shows a schematic view of the alleles. B6.Cg-E2f1Tg(Wnt1-cre)2Sor/J (Wnt1-Cre; Jackson Laboratory stock No. 022501) were crossed to mice carrying the *Sned1^fl^* allele to generate the NCC-specific *Sned1* knockout mouse line (*Sned1^NCC-^* allele).

The three mouse strains generated for this study have been deposited to Mutant Mouse Resource & Research Centers (MMRRC) repository: *Sned1^LacZ-Neo^* strain No. 065813, *Sned1^fl^* strain No. 065814, and *Sned1* strain No. 065815 and will all be made available upon publication of this manuscript.

Mice were subjected to blind visual genotyping and with a very high rate, knockout mice, whose genotype was confirmed by PCR (see below), were easily distinguished because of their craniofacial phenotype from wild-type and heterozygous littermates.

### Genotyping

DNA was extracted from tail samples from 14- or 21-day old pups, or from yolk sacs or tail samples from E18.5 embryos, according to standard procedures and genotyping was performed with the following primers: *Sned1 (WT allele: 513 bp; floxed allele: 614 bp):* Sned1_F TTCTTATTCACACCGTATGCCAGCC and Sned1_ttR CTAGTGGGCACTCATTCAGCAAACC; *Null allele of Sned1 (732 bp):* Sned1_F TTCTTATTCACACCGTATGCCAGCC and Sned1_R TCACATGAGCAACACGTTTGTAGG; *Neo cassette (603bp):* Neo_F GGGATCTCATGCTGGAGTTCTTCG and Sned1_ttR CTAGTGGGCACTCATTCAGCAAACC; *LoxP site after exon 2 (374 bp):* LoxP_F GAGATGGCGCAACGCAATTAATG and Sned1_R TCACATGAGCAACACGTTTGTAGGG; *Flp transgene (725 bp):* Flp_F CACTGATATTGTAAGTAGTTTGC and Flp_R CTAGTGCGAAGTAGTGATCAGG; *Cre transgene (700 bp):* Cre_F TGCTGTACTGGTTATGCGG and Cre_R TTGCCCCTGTTTCACTATCCAG.

PCR amplification was performed using the DreamTaq DNA polymerase (Thermo Fisher Scientific, Waltham, MA). Cycling conditions were: 95°C for 2 min.; 35 cycles at: 95°C for 30 sec., 60°C for 30 sec., 72°C for 45 sec.; final extension at 72°C for 3 min. Representative genotyping gels are presented in Figure 2B and Figure 7A.

### Survival analysis

Goodness-of-fit analysis was performed using a two-tailed chi-square (χ^2^) test (GraphPad). This statistical analysis compares expected (Mendelian distribution) and observed numbers of embryos (E18.5) or pups (p21) to determine whether survival correlates with animal genotypes.

### Isolation of mouse embryonic fibroblasts and quantitative PCR

Mouse embryonic fibroblasts (MEFs) were isolated from E13.5 to E15.5 embryos obtained by timed mating of heterozygous *Sned1^WT/LacZ-Neo^* mice. For quantitative PCR (qPCR), RNA was isolated from the cells using the RNeasy kit (Qiagen, Germantown, MD) and cDNA was synthesized by reverse transcription using the First-Strand cDNA Synthesis Kit (Promega, Madison, WI). qPCR reactions were performed using Bio-Rad SYBR Green Supermix (Bio-Rad, Hercules, CA) according to the manufacturer’s instructions. PCR conditions were 95°C for 10 minutes, followed by 40 cycles of 95°C for 20 seconds, 58°C for 30 seconds, and 72°C for 30 seconds. qPCR data analysis was performed using Bio-Rad CFX Manager Software. The following primers were used: Actin_F TGTATGAAGGCTTTGGTCTCC; Actin_R GTCTCAAGTCAGTGTACAGGC; Sned1_F GTAGATGGAAGAGGAAGAGTGAG; Sned1_R CTGTTCTTGGGTAGCTGGAG.

### β-galactosidase assay

β-galactosidase assay (LacZ staining) was performed on whole heterozygous *Sned1^WT/LacZ-Neo^* embryos (E11.5 and E13.5) according to previously published detailed protocols.^78,79^ For visualization of whole-mount staining, embryos were cleared by incubation in solutions of increasing glycerol and decreasing potassium hydroxide concentrations following a published protocol.^80^ LacZ-stained embryos were embedded in paraffin and sectioned, sections were then dewaxed and rehydrated, and hematoxylin and eosin counter-staining was performed following standard procedures.

### Microcomputed tomography image acquisition and analysis

An eXplore CT 120 μCT system (Northridge Tri-Modality Imaging Inc., Chatsworth, CA) was used for *in-vivo* imaging of a cohort of 19 adult (6-month-old) and 22 (p0.5) global *Sned1* knockout animals and control littermates. Mice were imaged under anesthesia (induced at 3% isoflurane in oxygen, maintained at between 2-2.5% during imaging) in groups of 4 in a custom mouse holder. Scanner settings were as follows: 720 views, 360-degree rotation, 70 kVp, 50 mA, 32 ms integration time with 2×2 detector pixel binning (isotropic nominal resolution of 50 microns). Data were reconstructed using the Parallax Innovations (Ilderton, ON, Canada) GPU accelerated reconstruction engine for the eXplore CT120.

A cohort of 31 adult (7 to 15-month-old) NCC-specific *Sned1* knockout animals and control littermates were scanned using a custom-built GE phoenix vļtomeļx s240 dual-tube X-ray μCT system (General Electric, Boston, MA) located at the University of Chicago. Data was acquired using a high power 240 kV tube. Scanner settings were as follows: 1,000 views, 360-degree rotation, 60 kVp, 350 μA, 250 ms integration time to achieve isotropic reconstructions with resolution (voxel size) of 40 microns. Data were reconstructed using the datosļx reconstruction software (General Electric, Boston, MA). The same parameter of beam hardening correction was applied to all samples. Reconstructed μCT scans of the mouse crania were converted to TIFF files and imported into 3D Slicer or ImageJ for further analysis.

### Measurement of long bone length

3D volume renderings of long bones were created from μCT scan files in 3D Slicer. Length of long bones was measured in mm. Unpaired t-tests were conducted to analyze differences in the length of long bones between genotype groups.

### Measurement of cranial suture length

Variations in the cranial sutures were visually observed in 3D Slicer.^81,82^ The nasal (anterior tip of nasal bones to premaxillary suture), frontal (premaxillary suture to coronal suture), and sagittal (coronal to interparietal suture) sutures were observed through the transverse plane and were measured antero-posteriorly (in mm), while the coronal sutures were observed through the sagittal plane. Unpaired t-tests were conducted to analyze differences in the length of the calvarial sutures between genotype groups.

### Assessment of nasal cavity symmetry and occlusion

Analyses were conducted using ImageJ software, using the “Multi VFF Opener” plug-in to open the μCT scan files. Variations in nasal cavity symmetry and occlusion were visually observed in 3D Slicer^81,82^ and scored.

### Landmarking and geometric morphometric analysis of the skull

We selected a total of 30 landmarks (22 traditional coordinate landmarks, 8 sliding semi-landmarks) which were deemed to represent well overall cranial morphology (*see positioning of landmarks in Supplementary Method 1*). The 8 sliding semi-landmarks were positioned along the nasal suture, as visual observation indicated that nasal and mid-facial morphology may be particularly affected in these mice. To undertake complex phenotyping, we used standard geometric morphometric (GM) methodologies for analyzing 3D shapes. First, a generalized Procrustes analysis (GPA)^83^ was run on the raw coordinate landmark data. This procedure scales, rotates, and translates landmark data to align them for further analysis. Importantly, this step (through scaling) reduces size to a separate variable so that shape can be studied in isolation.^84^ Semi-landmarks were slid, minimizing bending energy, as a part of the GPA. Next, to investigate shape variation in the cohort, we ran a principal component analysis (PCA) on the GPA data. Statistical significance of each of the first four PCs was assessed using a one-way non-parametric Kruskal-Wallis analysis of variance (ANOVA) by genotype followed by a Tukey’s HSD test.

### Landmarking and geometric morphometric analysis of the mandible

We selected a total of 18 traditional landmarks which were deemed to represent well overall mandibular morphology (*see positioning of landmarks in Supplementary Method 2*)^42^. Next, to investigate shape variation in the cohort, we ran a PCA on the GPA data. Statistical significance of each of the first four PCs was assessed using a one-way non-parametric Kruskal-Wallis ANOVA by genotype followed by a Tukey’s HSD test.

## Acknowledgements

The authors wish to acknowledge Dr. Aurora Burds Connor and Noranne Enzer from the ES Cell and Transgenics Facility (MIT) for their help generating the *Sned1* mouse lines, Lisa Billings and Katherine De La Hoz from the MIT Department of Comparative Medicine for their assistance with mouse husbandry, Kathleen Cormier from The Hope Babette Tang (1983) Histology Facility (MIT) for her help with preparation and sectioning of histological samples, Milton Cornwall-Brady and Dr. Scott Malstrom from the Animal Imaging & Preclinical Testing Core Facility (MIT), and April I. Neander (University of Chicago) for their assistance with μCT image acquisition. The authors also wish to acknowledge Dr. Patrick Murphy (Hynes lab) for noticing initially the craniofacial anomalies of the *Sned1* knockout mice, Ying Huang and Zhigang Jiang (Hynes lab) for assistance with genotyping, Martin Davis (Naba lab) for assistance with genotyping and mouse colony maintenance, and Ali Thahab (Naba lab) for his assistance with the analysis of human clinical data.

The *Sned1* mouse strains used for this research project were created from ES cell clone EPD0300_5_E02, obtained from the KOMP Repository (www.komp.org) and generated by the Wellcome Trust Sanger Institute. Targeting vectors used were generated by the Wellcome Trust Sanger Institute and the Children’s Hospital Oakland Research Institute as part of the Knockout Mouse Project (3U01HG004080).

## Supplementary Files

**Movies of the μCT scans used for occlusion and asymmetry assessment of nasal cavities.**

**Supplementary File 1.** File 1_WT_Adult.mp4

μCT scan of a 6-month-old wild-type (*Sned1^WT/WT^*) mouse from the global *Sned1* knockout transgenic line.

**Supplementary File 2.** File 2_Het_Adult.mp4

μCT scan of a 6-month-old heterozygous (*Sned1^WT/LacZ-Neo^*) mouse from the global *Sned1* knockout transgenic line.

**Supplementary File 3.** File 3_KO_Adult.mp4

μCT scan of a 6-month-old knockout (*Sned1^LacZ-Neo/LacZ-Neo^*) mouse from the global *Sned1* knockout transgenic line.

**Supplementary File 4.** File 4_WT_Neonate.mp4

μCT scan of a p0.5 wild-type (*Sned1^WT/WT^*) pup from the global *Sned1* knockout transgenic line.

**Supplementary File 5.** File 5_Het_Neonate.mp4

μCT scan of a p0.5 heterozygous (*Sned1^WT/LacZ-Neo^*) pup from the global *Sned1* knockout transgenic line.

**Supplementary File 6.** File 6_KO_Neonate.mp4

μCT scan of a p0.5 knockout (*Sned1^WT/LacZ-Nea^*) pup from the global *Sned1* knockout transgenic line.

**Supplementary File 7.** File 7_NCC_WTfl_Adult.mp4

μCT scan of a 9-month-old control (*Sned1^WT/fl^*) mouse from the NCC-specific *Sned1* knockout transgenic line.

**Supplementary File 8.** File 8_NCC_flfl_Adult.mp4

μCT scan of an 8-month-old control (*Sned1^fl/fl^*) mouse from the NCC-specific *Sned1* knockout transgenic line.

**Supplementary File 9.** File 9_NCC_Het_Adult.mp4

μCT scan of a 9-month-old heterozygous (*Sned1^WT/NCC-^*) mouse from the NCC-specific *Sned1* knockout transgenic line.

**Supplementary File 10.** File 10_NCC_KO_Adult.mp4

μCT scan of a 14-month-old knockout (*Sned^NCC-/NCC-^*) mouse from the NCC-specific *Sned1* knockout transgenic line.

**Supplementary Method 1.**
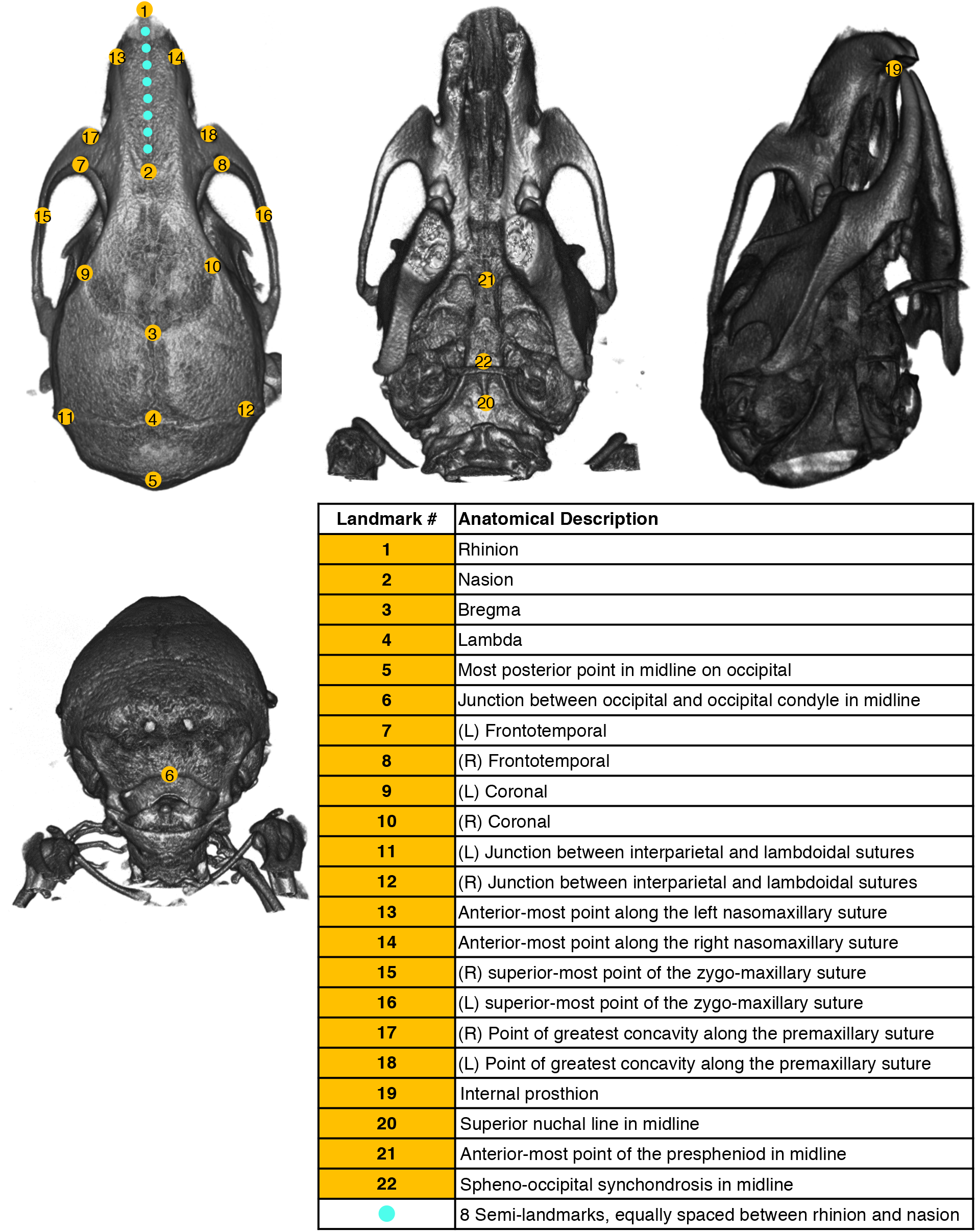
Diagram representing the positioning of calvarial landmarks.

**Supplementary Method 2.**
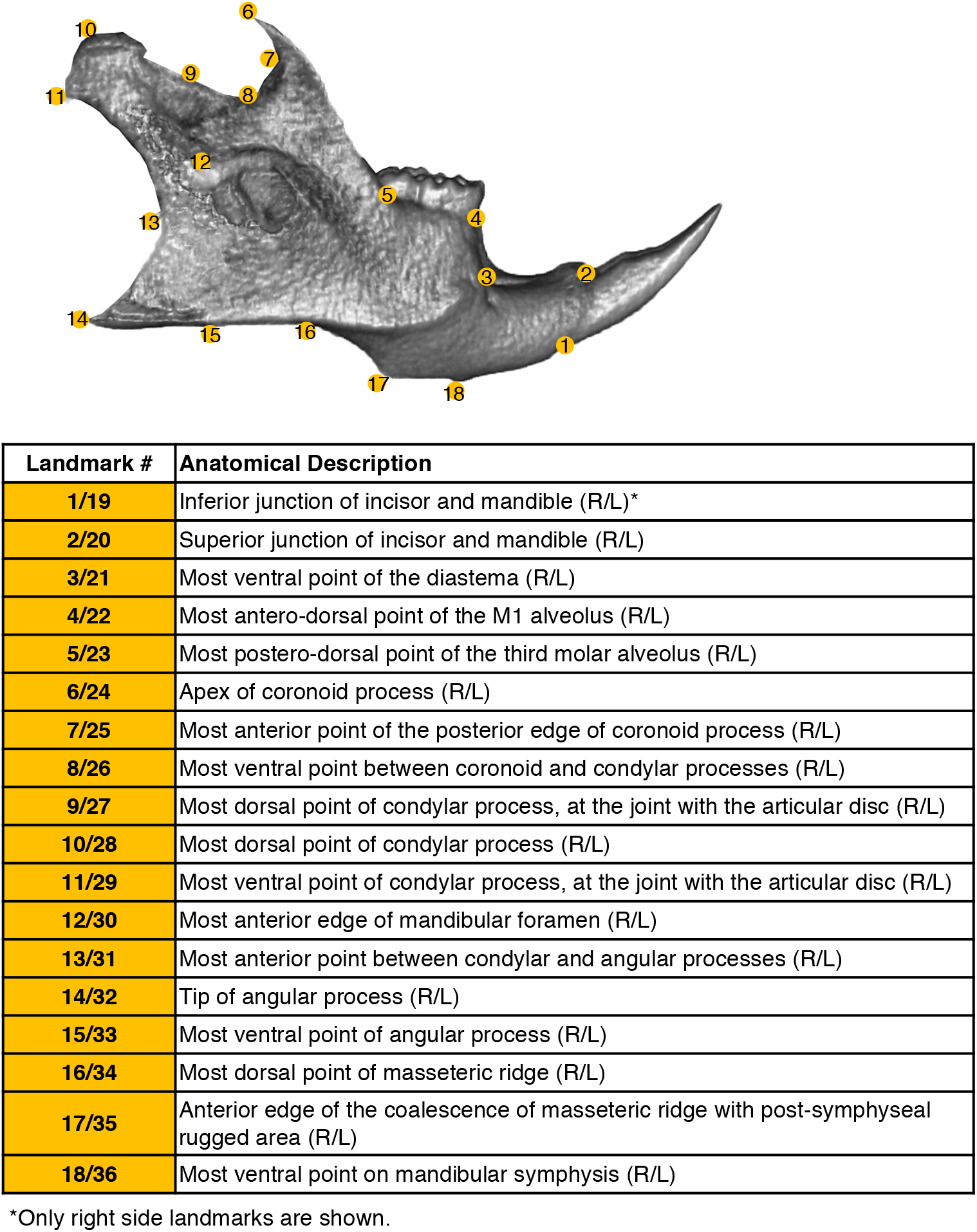
Diagram representing the positioning of mandibular landmarks.

